# Unexplained Repeated Pregnancy Loss is Associated with Altered Perceptual and Brain Responses to Men’s Body-Odor

**DOI:** 10.1101/2020.02.06.937029

**Authors:** Liron Rozenkrantz, Reut Weissgross, Tali Weiss, Inbal Ravrebi, Idan Frumin, Sagit Shushan, Lior Gorodisky, Netta Reshef, Yael Holzman, Liron Pinchover, Yaara Endevelt-Shapira, Eva Mishor, Edna Furman-Haran, Howard Carp, Noam Sobel

## Abstract

In the *Bruce effect*, pregnant mice remember the odor of the fathering male, and miscarry in response to the odor of a male stranger. Humans experience a high rate of unexplained spontaneous miscarriage. Could it be that a portion of these miscarriages reflect a Bruce-like effect? Given ethical constraints on a direct test, we instead probed for circumstantial evidence in women with repeated pregnancy loss (RPL). Consistent with a Bruce-like effect, women with RPL remembered the body-odor of their spouse, but controls could not. Also consistent with a Bruce-like effect, body-odor from a stranger man caused increased activity in the hypothalamus of women experiencing RPL, yet decreased activity in the hypothalamus of women controls. Finally, RPL was associated with reduced olfactory-bulb volume. Although not causal, these observations link RPL with an altered behavioral and brain response to men’s body-odor, implicating the olfactory system in this poorly understood or managed condition.

## Introduction

A remarkable ~50% of all human conceptions, and ~15% of documented human pregnancies, end in spontaneous miscarriage (Rai and Regan, 2006). Only a portion of these miscarriages, however, can be explained (Clifford et al., 1994; Stephenson, 1996; Jaslow et al., 2010), suggesting the presence of major yet-unidentified human-miscarriage-causing factors. Pregnant mice at an early phase of pregnancy miscarry following exposure to bodily odors emitted from a male who did not father the pregnancy (Bruce, 1959). This effect, named the *Bruce effect* after Hilda Margaret Bruce who discovered it, is highly robust, occurring in ~70-80% of exposures (Bruce, 1959, 1960). Although the *Bruce effect* was initially characterized as pregnancy block prior to embryo implantation in mice (Bruce, 1959, 1960; Bruce and Parrott, 1960), it has since been established in several additional rodent species, and at various stages of pregnancy, including post-implantation, mid-gestational and up to day-17 of a 23-day pregnancy (Eleftheriou et al., 1962; Clulow and Langford, 1971; Mallory and Brooks, 1980; Mahady and Wolff, 2002). Bruce-like effects have also been hypothesized, although not verified by manipulation, in lions (Bertram, 1975; Packer and Pusey, 1983), wild horses (Berger, 1983), dogs (Bartos et al., 2016), and primates (Pereira, 1983; Mori and Dunbar, 1985; Agoramoorthy et al., 1988; Colmenares and Gomendio, 1988). Of the latter, a particularly comprehensive study observed pregnancy termination in ~80% of gelada baboon females following replacement of the dominant male (Roberts et al., 2012). Given these examples, we hypothesized that the olfactory system, and particularly responses to men’s body-odor, may be involved in human spontaneous miscarriage as well.

Two alternative *Bruce effect* mechanisms have been proposed. In the first, male-specific urinary pheromones are transduced at the female’s vomeronasal organ (VNO) (Zufall and Leinders-Zufall, 2007; Becker and Hurst, 2008). These chemical signals trigger a hypothalamic-dependent downstream neuroendocrine response that includes an increase in dopamine and reduction in prolactin, a hormone crucial for maintenance of the curpus luteum, and thus for embryo implantation (Rosser et al., 1989; Becker and Hurst, 2008; Brennan, 2009). However, if the female learns and remembers the individual-specific pheromones during mating, subsequent encounters with this chemical signature will release noradrenaline, inhibiting the VNO-dependent cascade, thus preventing pregnancy disruption (Brennan and Zufall, 2006; Becker and Hurst, 2008). In support of this model, removing the female VNO significantly reduces or prevents pregnancy block (Bruce, 1960; Bellringer et al., 1980; Lloyd-Thomas and Keverne, 1982). An alternative *Bruce effect* mechanism involves estradiol (E2), a metabolic product of testosterone. By this model, when a female is exposed to a male’s urine, E2 enters the bloodstream via nasal ingestion, travels to the uterus, and disrupts implantation (Valbuena et al., 2001; Ma et al., 2003; Guzzo et al., 2010). In support of this model, the body-odor of castrated males fails to terminate pregnancy, but this is reversed upon administration of testosterone (Bruce, 1965; Guzzo et al., 2010). A limitation of both models is that they account for pregnancy block at pre-implantation alone, yet as noted, Bruce-like effects are evident later in pregnancy as well (Eleftheriou et al., 1962; Clulow and Langford, 1971; Mallory and Brooks, 1980; Mahady and Wolff, 2002).

Testing the hypothesis behind the *Bruce effect* in laboratory animals was straight-forward: researchers systematically exposed pregnant females to stranger male body-odor in an effort to cause pregnancy-block or miscarriage. In humans, however, ethical considerations prevent us from engaging in such an experiment. We obviously cannot try to cause human miscarriage. The only interventional experiment and potentially causal test of our hypothesis that we can think of, entails separating miscarriage-prone women from the company of men during early pregnancy (by removing either the men or the women), and then estimating whether this protected the pregnancy based on outcome compared to past individual experience. For example, current participant RPL36 experienced 12 consecutive miscarriages (Supplementary Table 1). If upon separation from the company of men she would now maintain a pregnancy, this could be seen as telling. Coincidently, we note that in several tribal societies it is indeed customary in early pregnancy to separate women from the company of men (Van Gennep, 1960). Moreover, human repeated pregnancy loss (RPL) has been associated with a specific polymorphism in an olfactory receptor gene (OR4C16G>A) (Ryu et al., 2019). Nevertheless, this scant anecdotal evidence is not reason enough to test a socially drastic manipulation such as separation, especially during an emotionally complicated time. With that in mind, we set out to probe for additional circumstantial evidence that may link between human miscarriage and men’s body-odor. About 2-4% of couples that experienced a spontaneous miscarriage (i.e., ~1% of women) are prone to RPL, namely two or more consecutive miscarriages that mostly remain unexplained despite extensive medical investigation (Clifford et al., 1994; Stephenson, 1996; Jaslow et al., 2010). Here we asked whether women who experience RPL have a different perceptual and brain response to men’s body-odor.

## Results

### Whereas control women cannot identify their spouse by smell, women with RPL can

The *Bruce effect* mechanism requires the female to remember and identify the body-odor of the fathering male (Leinders-Zufall et al., 2004; Brennan and Zufall, 2006). To characterize olfactory spouse-recognition in humans we tested 33 women with repeated pregnancy loss (RPL) (mean age = 34.7 ± 6.4, mean number of miscarriages = 4.2) and 33 matched controls (mean age = 35.6 ± 4.1, never experienced miscarriage). We used a three-alternative forced-choice (3AFC) paradigm where on each trial the participant was asked to identify her spouse from three alternatives (delivered in specially-designed body-odor sniff-jars, see Supplementary Fig. 1); one containing the body-odor of her spouse, and two containing distractor odors; one from a non-spouse man (*Stranger*) and one *Blank* (carrier alone). Previous results on spouse odor identification in the general population are mixed, with two studies reporting ~33% group-level performance (Hold and Schleidt, 1977; Mahmut et al., 2019) but one study reporting ~74% performance (Lundström and Jones-Gotman, 2009). Here, consistent with the two former studies, as a group, control women were unable to identify their spouse above chance levels (chance = 33.3%, mean control = 28% ± 34%, Shapiro-Wilk test of normality (SW) = 0.77, p < 0.001, difference from chance: Wilcoxon W = 206, p = 0.18, effect size estimated by rank biserial correlation (RBC) = 1.17 (parametric comparison: t(32) = 0.88, p = 0.39, Cohen’s d = 0.15)). In contrast, remarkably, RPL women were able to identify their spouse by body-odor alone, at a level that was both significantly higher than expected by chance (chance = 33.3%, mean RPL = 57% ± 41%, SW = 0.81, p < 0.001, difference from chance: Wilcoxon W = 440, p = 0.004, RBC = 1.71 (parametric comparison: t(32) = 3.32, p = 0.002, Cohen’s d = 0.58), and two-fold that of controls (Mann-Whitney U = 332.5, p = 0.005, RBC = 0.39, (parametric comparison: t(64) = 3.12, p = 0.003, Cohen’s d = 0.77) (Fig. 1a). In other words, consistent with the mechanistic underpinnings of the Bruce effect in mice, an increased tendency for miscarriage in women is associated with better identification of spouse body-odor.

**Fig. 1.**
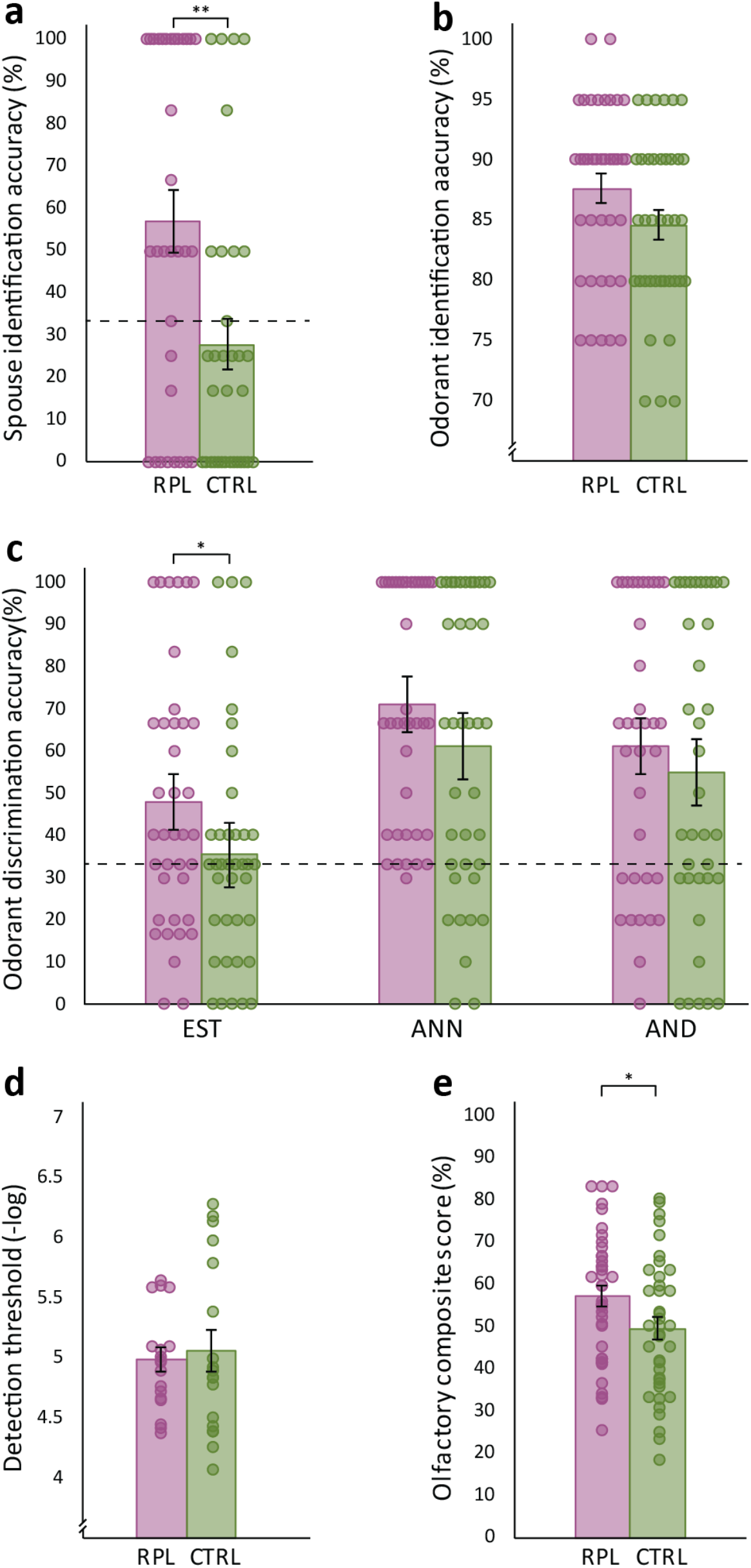
Women with RPL can identify their spouse by smell. RPL (purple) and control (green) women were tested for various olfactory facets: **a.** Spouse identification (n=66). **b.** Every-day odorant identification (n=76). **c.** Monomolecule discrimination (EST, ANN, AND) (n=76). **d.** DMTS threshold (n=36). **e.** A composite score of identification, discrimination and threshold (n=78). Bar graphs depict means, each dot represents a participant, dots are jittered to prevent overlay, error bars are s.e.m. *p<0.05, **p<0.01. Black dashed line indicates chance level.

### Women with RPL have otherwise only marginally better olfaction than controls

In order to determine whether this advantage at identifying spouse body-odor was a reflection of a generally better sense of smell, we tested olfactory performance at several additional tasks. To test ordinary odor identification, 38 women with RPL and 38 matched controls conducted a four-alternative forced-choice identification test using 20 every-day odorants such as *soap* and *peanuts*. We observed only a trend towards better identification in RPL women than in controls (mean RPL = 87.6% ± 7.2, Controls =84.6% ± 7.3, RPL SW = 0.91, p = 0.005, control SW = 0.92, p = 0.009, Mann-Whitney U = 560.5, p = 0.087, RBC = 0.22 (parametric comparison: t(74) = 1.82, p = 0.073, Cohen’s d = 0.42)) (Fig. 1b). Subsequently, 38 women with RPL and 38 controls were tested for detection of three odorant monomolecules that have often been studied in the context of human social chemosignaling(Jacob and McClintock, 2000). The monomolecules were Androstenone (ANN), Androstadienone (AND), and Estratetraenol (EST). We used each odorant at a fixed concentration, and tested its discrimination from blank using a 3AFC test. To account for the non-normal distribution of the data, a linear mixed-model with fixed effects of Group and Odor, with random intercept per participant and a random slope for odor within participant, revealed a main effect of Odorant (F(2,143) = 17.1, p < 0. 0001), a modest effect of Group (F(1,74) = 3.97, p = 0.05) and no interaction (F(2,143) = 0.27, p = 0.77) (for comparison, a parametric repeated-measures analysis of variance (ANOVA) with factors of Group (RPL/Control) and Odorant (ANN/AND/EST) revealed a main effect of Odorant (F(2,148) = 17.5, p = 1.5e^−7^), but only a trend toward a main effect of Group (F(1,74) = 3.28, p = 0.074), and no interaction (F(2,148) = 0.31, p = 0.74) (Fig. 1c). The main effect of Odorant reflected that at these particular concentrations, EST was harder to detect than both AND and ANN (mean EST = 41.8 ± 29.4, mean AND = 58.1 ± 34.6, mean ANN = 66.2 ± 31: AND vs. ANN Wilcoxon W = 858, p = 0.068, RBC = 0.29 (parametric comparison: t(70) = 1.7, p = 0.093, Cohen’s d = 0.2); AND vs EST Wilcoxon W = 451, p = 6.4e^−4^, RBC = 0.51 (parametric comparison: t(70) = 3.5, p = 8.2e^−4^, Cohen’s d = 0.42); ANN vs. EST: Wilcoxon W = 237, p = 6e^−7^, RBC = 0.74 (parametric comparison: t(75) = 5.86, p = 1.2e^−7^, Cohen’s d = 0.67) (Fig. 1c). The trend towards a main effect of Group reflected that RPL participants again performed slightly better than controls, but this effect was near-significant for the odorant EST alone (EST: RPL mean = 48.1% ± 30.3%, Control mean = 35.4% ± 27.4%, Mann-Whitney U = 544.5, p = 0.065, RBC = 0.25 (parametric comparison: t(74) = 1.9, p = 0.06, Cohen’s d = 0.44); AND: RPL mean = 61.1% ± 32.5%, Control mean = 55.1% ± 36.8%, Mann-Whitney U = 585, p = 0.6, RBC = 0.07 (parametric comparison: t(69) = 0.73, p = 0.47, Cohen’s d = 0.17); ANN: RPL mean = 71.1% ± 27.1%, Control mean = 61.3% ± 34%, Mann-Whitney U = 587.5, p = 0.15, RBC = 0.19 (parametric comparison: t(74) = 1.38, p = 0.17, Cohen’s d = 0.32) (Fig. 1c). Finally, a group of 18 women with RPL and 18 controls were also available for a lengthy 7-reversal staircase paradigm using the alliaceous odorant dimethyl trisulfide (DMTS). We observed no significant difference between controls and RPL (RPL threshold, minus log concentration: 4.98 ± 0.41, control threshold: 5.06 ± 0.72, RPL SW = 0.92, p = 0.13, control SW = 0.9, p = 0.052, t(34) = 0.42, p = 0.68, Cohen’s d = 0.14) (Fig. 1e). In reviewing the above results at olfactory identification, discrimination, and detection, women with RPL repeatedly scored slightly better than controls, but this difference was not significant. For added power and sensitivity, in a final comparison we combined the above results, generating a composite olfaction score for each participant (reflecting her available scores at identification, discrimination, and detection) ranging between 0 and 100 (see Methods). This enabled a comparison between 39 women with RPL and 39 controls. We observed a small but significant difference between the two groups (mean composite RPL = 57.13% ± 15.23, Controls = 49.46% ± 16.14, RPL SW = 0.97, p = 0.49, control SW = 0.98, p = 0.67, independent t-test t(76) = 2.16, p = 0.034, Cohen’s d = 0.49). We further tested for any relationship between this composite score and the overwhelmingly better olfactory identification of spouse in RPL versus controls. We observed no correlation across all participants, but a trend in RPL (RPL group (n = 33): Spearman Rho = −0.33, p = 0.06; Control group (n = 29): R = 0.06, p = 0.76; Both groups together: R = −0.02, p = 0.88). In conclusion, women with RPL were two-fold better than control women at identifying their spouse by smell, and marginally but significantly better than control women at ordinary olfactory tasks.

### Women with RPL have altered perception of men’s body-odor

Given that the advantage at identifying spouse by smell was unrelated to general olfactory abilities, we next asked whether it was reflected in any explicit perceptual olfactory attributes in body-odor. A cohort of 18 women with RPL and 18 controls rated men’s body-odors on four traits: Intensity; Pleasantness; Sexual attraction; and Fertility. Each woman blindly (but see later note) rated stimuli of three kinds: *Blank* (carrier alone), *Stranger* (a non-spouse man), and their actual *Spouse*. To account for the non-normal distribution of the data, a linear mixed-model with fixed effects of Group, Odor and Descriptor, and random effects of participant, revealed a main effect of Descriptor (F(3,374) = 9.67, p =3.69e^−6^), and significant interactions of Odor x Descriptor (F(6,374) = 5.69, p =1.13e^−5^) and Group x Odor (F(2,374) = 7.77, p = 4.95e^−4^) (parametric comparison: a multivariate repeated measures ANOVA with factors of Group (RPL/Control), Odor (Blank/Stranger/Spouse) and Descriptor (Intensity/Pleasantness/Sexual attraction/Fertility) revealed a main effect of Descriptor (F(3,102) = 12.4, p = 5.6e^−7^), a significant interaction of Odor x Descriptor (F(6,204) = 8.27, p = 5.1e^−8^) and a significant interaction of Odor x Group (F(2,68) = 3.43, p = 0.038) (Figure 2). The main effect of Descriptor is uninteresting in that it merely reflected that descriptors were applied differently (mean Intensity: 0.49 ± 0.21, mean Pleasantness: 0.42 ± 0.14, mean Sexual attraction: 0.33 ± 0.15, mean Fertility: 0.38 ± 0.15; Wilcoxon pairwise comparisons except intensity-pleasantness: all W > 475, all p < 0.026, intensity-pleasantness Wilcoxon W = 431, p = 0.127 (parametric comparison, all t(35) > 2.12, all p < 0.041, all Cohen’s d > 0.35) (Supplementary Figure 2). The Odor x Descriptor interaction was carried by intensity ratings alone (F(2,68) = 16.2, p =1.72e^−6^; all other descriptors: all F(2,68) < 2.05, all p > 0.13). This intensity difference was carried solely by differences from Blank, which was, unsurprisingly, less intense than all other otherwise (and importantly) equally-intense stimuli (mean Intensity ratings: Blank: = 0.31 ± 0.26, Stranger = 0.58 ± 0.23, Spouse: = 0.59 ± 0.35. Blank vs. Stranger and Spouse: both Wilcoxon W < 98, p < 0.0001, RBC > 0.7 (parametric comparison: Both t(35) > 4.7, both p < 3.8e^−5^, both Cohen’s d > 0.77); Stranger vs Spouse: Wilcoxon W < 319, p = 0.83, RBC = 0.04 (parametric comparison: t(35) = 0.15, p = 0.88, Cohen’s d = 0.03) (Supplementary Figure 2). As to the Odor x Group interaction, whereas RPL and Control women rated Blank and Spouse the same (RPL Blank: = 0.36 ± 0.2; Control Blank: = 0.39 ± 0.18, Mann-Whitney U = 173, p = 0.74, RBC = 0.07 (parametric comparison: t(34) = 0.49, p = 0.63, Cohen’s d = 0.16); RPL Spouse: = 0.46 ± 0.23; Control Spouse: = 0.37 ± 0.21, Mann-Whitney U = 121, p = 0.2, RBC = 0.25 (parametric comparison: t(34) = 1.25, p = 0.22, Cohen’s d = 0.42), Stranger was rated lower by RPL than by Controls (RPL = 0.36 ± 0.18; Controls = 0.49 ± 0.14, Mann-Whitney U = 244, p = 0.009, RBC = 0.51(parametric comparison: t(34) = 2.35, p = 0.025, Cohen’s d = 0.78) (Fig. 2). This effect that materialized in the combined descriptors was mostly carried by a powerful difference is rated fertility attributed to the body-odors (Supplementary Figure 2). In the spouse condition, RPL and control women smelled different stimuli (each woman smelled her own spouse), so this was not an optimal condition for estimating differences in olfactory perception across these groups (we acknowledge that this condition adds the tantalizing possibility that in addition to altered olfaction in women with RPL, men in RPL relationships may have a unique body-odor, yet we do not have statistical power for this separate question within the current study). However, in the *stranger* condition, both groups are smelling common stimuli, and we observed a significant difference in perception. In other words, we observed a difference in the overall explicit perceptual ratings applied to men’s body-odor by women with RPL versus controls.

**Fig. 2.**
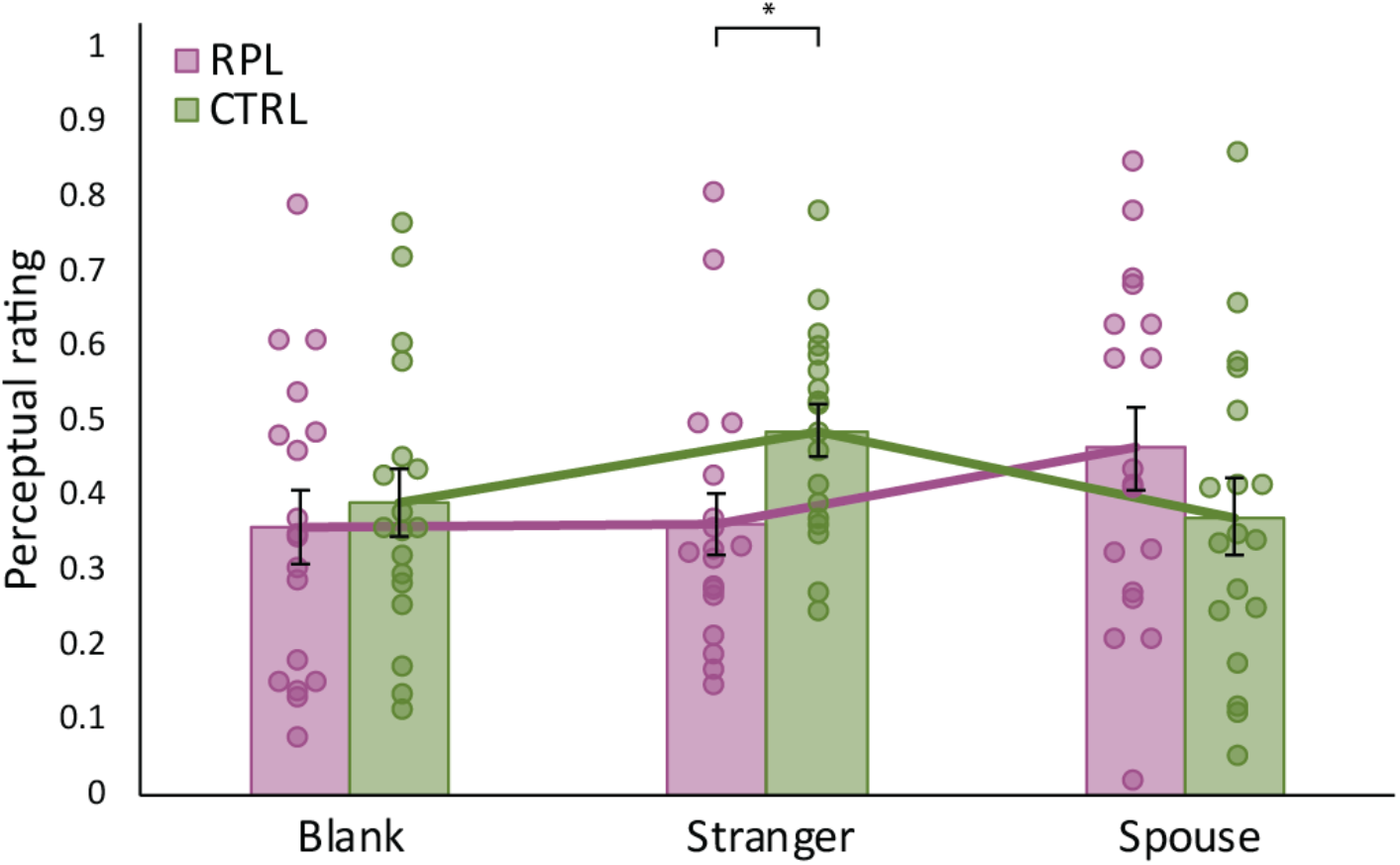
Women with RPL have altered perception of men’s body-odor. RPL (purple) and control (green) women ratings of stranger male body-odor, score combines ratings of pleasantness, sexual attraction, intensity and fertility attributed to the odor (see separate ratings in Supplementary Figure 2). n=36. Bar graphs depict means, each dot represents a participant, dots are jittered to prevent overlay, error bars are s.e.m. *p<0.05.

### Women with RPL have smaller olfactory bulbs and shallower olfactory sulci

Results to this point imply an altered sense of smell in women with RPL, specifically in relation to men’s body-odor. To probe for brain correlates of this difference, we used magnetic resonance imaging (MRI) to scan 23 women with RPL and 23 controls. To compare brain structure, we applied both a targeted and a hypothesis-free approach. In the targeted investigation we compared brain volume in the two primary regions of interest where brain structure has previously been related to olfactory function, namely the olfactory bulbs (Buschhüter et al., 2008) and the olfactory sulci (Rombaux et al., 2010). We observed that women with RPL have significantly smaller olfactory bulbs (all SW > 0.94, all p > 0.2; Right bulb volume: RPL: 45.9 ± 12 mm^3^, control: 55.4 ± 9.7 mm^3^, t(44) = 2.96, p = 0.005, Cohen’s d = 0.87; Left bulb volume: RPL: 46.4 ± 8.9 mm^3^, control: 55.7 ± 9.4 mm^3^, t(44) = 3.45, p = 0.001, Cohen’s d = 1.02; Average bulb volume: RPL: 46.1 ± 9.9 mm^3^, control: 55.5 ± 9 mm^3^, t_44_ = 3.37, p = 0.0016, Cohen’s d = 0.99) (Fig. 3a), and significantly shallower olfactory sulci (Right olfactory sulcus depth did not reach significance: RPL: 6.4 ± 1.1 mm, control: 7.1 ± 1.4 mm, t(44) = 1.68, p = 0.1, Cohen’s d = 0.5; Left olfactory sulcus depth: RPL: 6.1 ± 1.4 mm, control: 7 ± 1.4 mm, t(44) = 2.22, p = 0.03, Cohen’s d = 0.66; Average olfactory sulcus depth: RPL: 6.3 ± 1.1 mm, control: 7 ± 1.3 mm, t_44_ = 2.12, p = 0.04, Cohen’s d = 0.62) (Fig. 3b). To verify that this difference was not merely a reflection of smaller heads/brains, we conducted the same analysis on Total Intracranial Volume (TIV), and observe no difference between RPL women and controls (TIV: RPL mean: 1366 ± 134 cm^3^, Control mean: 1390 ± 93 cm^3^, RPL SW = 0.95, p = 0.31, control SW = 0.94, p = 0.2, t(43) = 0.69, p = 0.5, Cohen’s d = 0.21). As olfactory bulb volume has been associated with olfactory performance (Buschhüter et al., 2008; Rombaux et al., 2010), we tested for such correlations in these data. The one olfactory test that was completed by all 46 scanned participants was the odorant identification test, which revealed no correlation with bulb volume (RPL: n=23, Spearman Rho = 0.25, p = 0.25. Control: n=23, Spearman Rho = 0.146, p = 0.507. Combined: n=46, Spearman Rho = 0.068, p = 0.652). To further probe this issue we assigned an olfaction composite score to each participant, reflecting all the olfactory tasks she completed (see Methods). Using this more comprehensive olfactory score, we observed a significant correlation in the RPL cohort alone (RPL: n=22, Spearman Rho = 0.45, p = 0.037; Control: n=23, Spearman Rho = −0.055, p = 0.8; Combined: n=45, Spearman Rho = 0.065, p = 0.67).

**Fig. 3.**
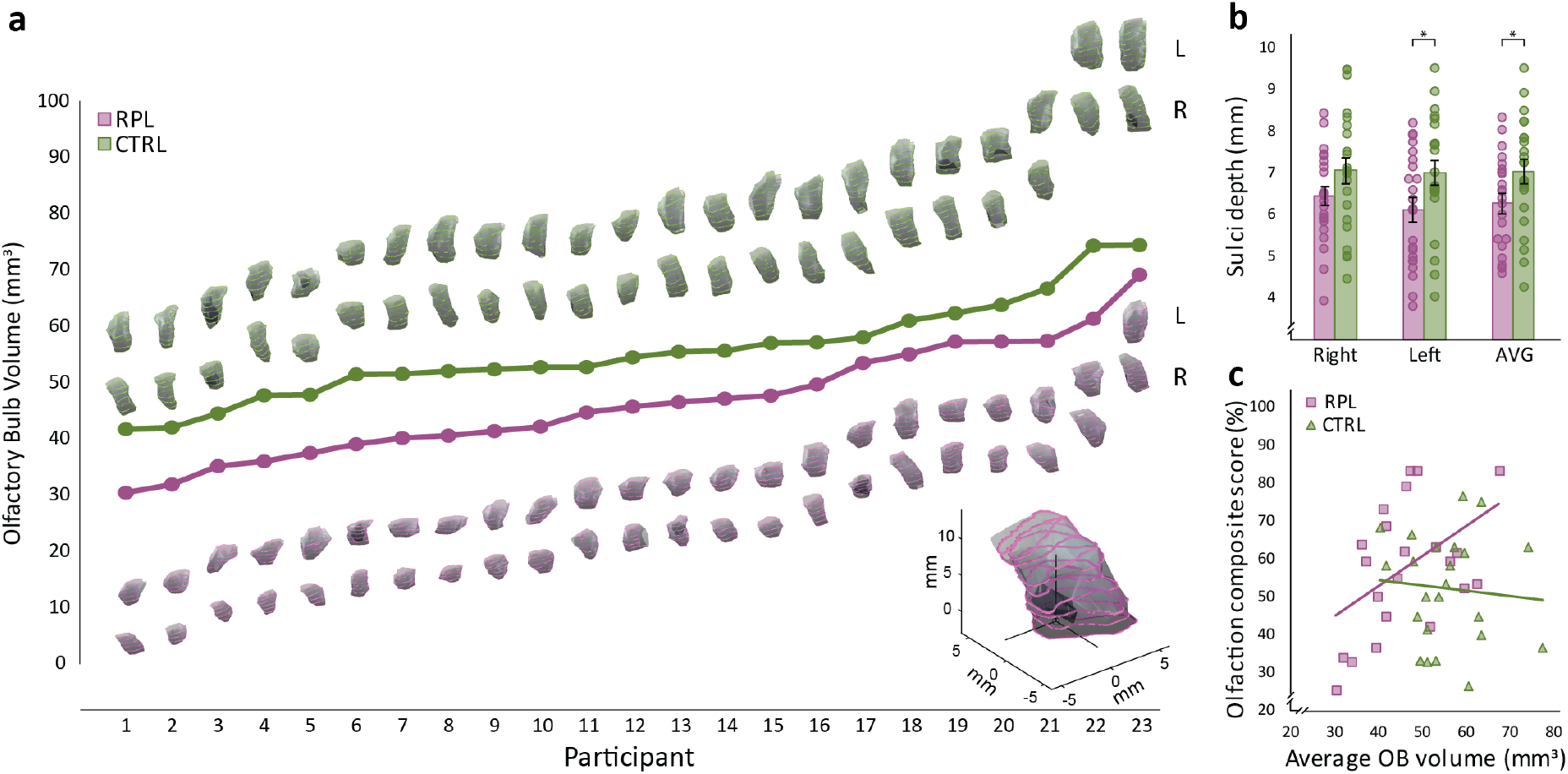
Women with RPL have smaller olfactory bulbs and shallower olfactory sulci. **a.** Olfactory bulb volume. A 3D reconstruction of RPL (purple) and control (green) participants’ left (upper row of the two) and right (bottom row) olfactory bulbs. Bulbs sorted by size (see Supplementary Figure 3 for sort by participant (RPL to Control) match). **b.** Olfactory sulci depth (right, left and average) of RPL (purple bars) and control (green bars) participants. Bar graphs depict means, each dot represents a participant, dots are jittered to prevent overlay. **c.** The relation between olfactory composite scores and olfactory bulb volumes for RPL (purple) and control (green) participants. n=46. Each square (RPL) and triangle (control) represent a participant. error bars are s.e.m. *p<0.05.

Next, in a hypothesis-free analysis we conducted whole-brain voxel-based morphometry (VBM). This analysis, with correction for the multiple comparisons associated with the entire brain, uncovered no significant group differences. We note that if we repeat this analysis at strict threshold (p < 0.0001) but without correction for multiple comparisons, it implied reduced gray matter in the right fusiform in RPL (Supplementary Figure 4). Hence, structural brain imaging uncovered a significant difference in the olfactory structures (olfactory bulb and olfactory sulci) of women who experience RPL, and hinted at a possible difference in non-olfactory structures (right fusiform) that have been implicated in social chemosignaling (Zhou and Chen, 2008). We next turn to functional imaging.

### Women with RPL have an altered brain response to men’s body-odor

We used functional magnetic resonance imaging (fMRI) in 23 women with RPL and 23 controls who watched varying-arousal movie-clips during subliminal exposure to body-odor from a *Stranger*. We chose not to use spouse body-odors in this particular experiment because this may have presented a confound: As noted, women with RPL can recognize their spouse’s body-odor. This pronounced difference between cohorts was overwhelmingly apparent in the previous rating experiment, where although blind to condition, upon presentation with spouse body-odor, several women from the RPL cohort spontaneously said “oh, that is my spouse”. This never happened once in the control cohort. Thus, if we used spouse odors in the MRI, we would risk the confound of only RPL women knowing that any odors, let alone their spouse body-odors, were presented. Moreover, given that RPL relationships may involve significant emotional strain (Kolte et al., 2015), this potential confound would be exacerbated. Finally, the Bruce effect, namely the conceptual framework of our investigation, is driven by *stranger* males. Thus, we chose to use male body-odor collected from donors unrelated to all study participants, and use the odor at subliminal levels that were not spontaneously detected.

The functional study was made of two consecutive scans, one with olfactometer-delivered undetected stranger body-odor, and one with olfactometer-delivered blank (counterbalanced for order across participants). In each such scan each participant observed 40 12s-long movie clips, and after each, participants self-rated the level of emotional arousal the clip induced (using an 8-level response box). Like in the structural analysis, in the functional analysis we also applied both a targeted and hypothesis-free approach. In the targeted approach we first concentrated on the hypothalamus, the primary brain structure implied in mediating the Bruce effect (Rosser et al., 1989; Yoon et al., 2005; Brennan, 2009). The hypothalamus has also been indicated in women’s responses to a molecule (AND) that may occur in men’s body-odor (Savic et al., 2001). We used a hypothalamic region of interest (ROI) independently defined in Neurosynth (Yarkoni et al., 2011), and applied a contrast of Group (RPL/Control) and Stimulus (Undetected body-odor/Blank) with parametric modulation for the self-rated level of emotional arousal. Following small-volume correction for multiple comparisons, we observed a remarkable effect in the hypothalamus (Z > 2.3, Cluster-corrected) (Fig. 4a). Although the p-value associated with this result is the statistical parametric map statistic (p < 0.01, corrected), we used a repeated-measures ANOVA applied to the parameter estimates only to understand the directional drivers of this effect. We observed that whereas body-odor increased hypothalamic activation in RPL (t(22) = 2.81, p = 0.01), it decreased hypothalamic activation in controls (t(22) = 2.41, p = 0.025), and the interaction was powerful (F(1,44) =13.4, p = 0.0007) (Fig. 4a). In turn, a hypothesis-free approach full-brain contrast ANOVA with correction for the multiple comparisons involved, did not uncover additional brain regions differently impacted by arousal as a function of exposure to body-odor in RPL vs. controls.

**Fig. 4.**
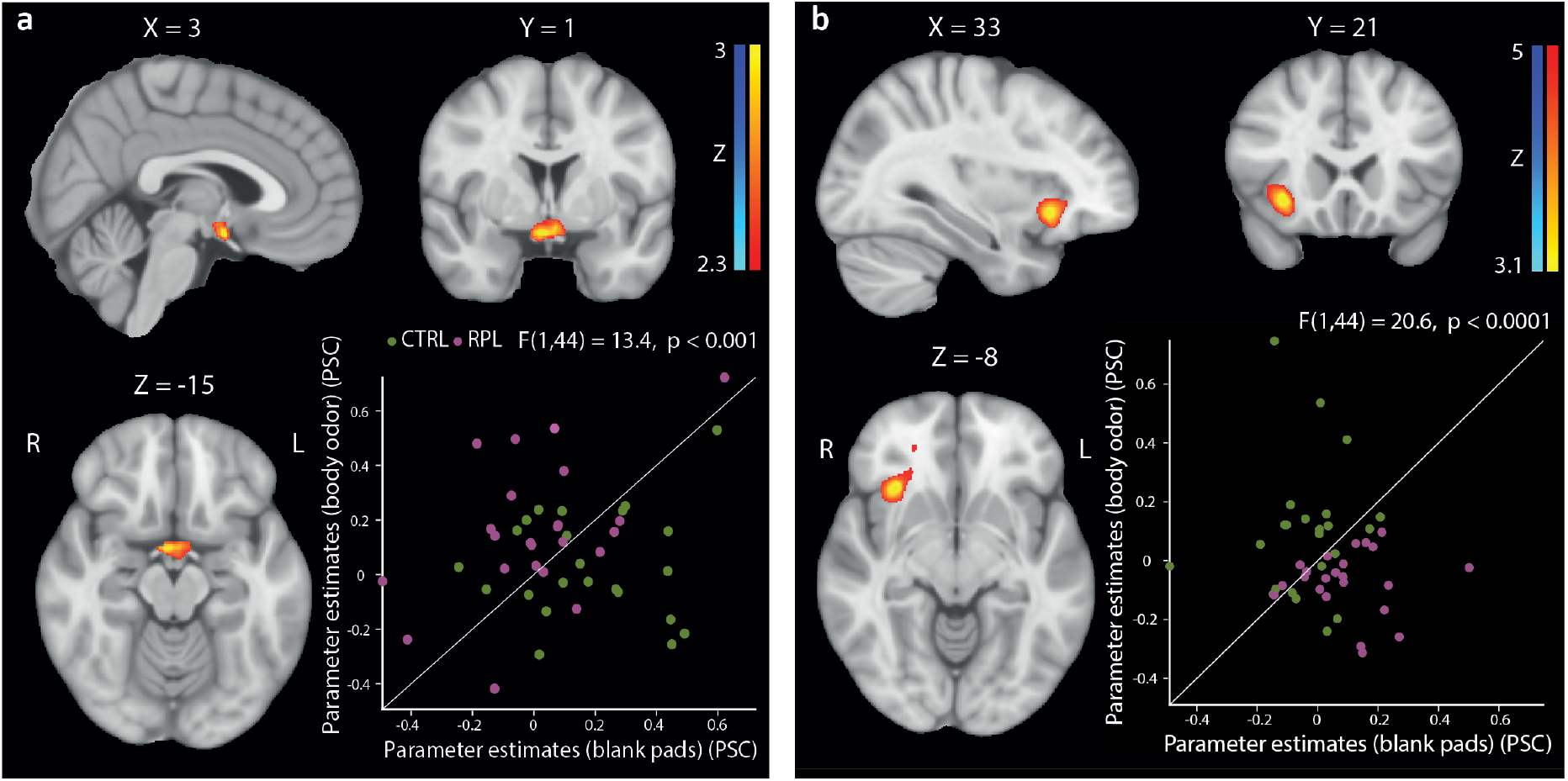
Women with RPL have an altered brain response to male body-odor. **a.** Hypothalamus blood-oxygen level-dependent (BOLD) activity in RPL (stranger > blank), compared to control. **b.** Whole brain PPI test, reflecting greater correlation with hypothalamus (seed ROI) time series for all emotionally-weighted-movie-clips > fixation. Both scatterplots reflect the parameter estimates (PE) of each participant (RPL in purple, controls in green): dots above the slop line represent higher PE for *Stranger* body odor, and dots below the line represent higher PE for *Blank*. n=46. All coordinates in MNI. PSC=percent signal change

Given the hypothalamic group-difference, we next used psychophysiological interaction (PPI) analysis (Friston et al., 1997) to ask whether functional connectivity with this region differed across groups as a function of exposure to body-odor. We observed that with increased arousal, body-odor increased connectivity between the hypothalamus and right insula significantly more in controls than in RPL (peak Montreal Neurological Institute coordinates (MNI): [33,21,-8], 2336 voxels, Cluster-corrected z > 3.1) (Fig. 4b) We again note that the p-value associated with this finding is the above PPI mapping statistic (p < 0.001, corrected), but to only verify the directionality of the effect we applied a repeated-measures ANOVA on the parameter estimates, revealing that the decrease was significant in RPL (t(22) = 2.45, p = 0.022), the increase was significant in Controls (t(22) = 4.41, p = 0.0002), and the interaction was powerful (F(1,44) = 20.6, p = 0.00004) (Figure 4b). Patterns of brain activity can be impacted by anxiety (Holzschneider and Mulert, 2011), which can be increased in RPL (Mevorach-Zussman et al., 2012). To address this possible source of variance in our data, all participants completed the trait-anxiety questionnaire (State-Trait Anxiety Inventory; STAI (Teichman and Melnick, 1979)), and a personality questionnaire (The “Big Five” Inventory (Etzion and Laski, 1998)). We observed that both RPL and controls had moderate anxiety levels, and there was no significant difference between the groups (STAI: RPL = 39.5 ± 8, Control = 39.87 ± 10.86, SW = 0.97, p = 0.3, t(44) = 0.14, p = 0.89, Cohen’s d = 0.041). Moreover, we observed no significant differences in personality (Extroversion (E), Conscientiousness (C), Neuroticism (N) Agreeableness (A), Openness to experience (O)); RPL: E = 3.67 ± 0.59, C = 3.71 ± 0.49, N = 3.05 ± 0.67, A = 3.87 ± 0.46, O = 3.58 ± 0.67. Control: E = 3.51 ± 0.64, C = 3.94 ± 0.41, N = 2.97 ± 0.65, A = 3.73 ± 0.64, O = 3.73 ± 0.44. All SW > 0.957, all p > 0.092, all t(44) < 1.68, all p > 0.1, all Cohen’s d < 0.5). Therefore, the unique pattern of brain activity in response to body-odor in RPL was not a reflection of differences in anxiety or personality across cohorts.

Finally, given these perceptual and brain differences, one may postulate a generalized different impact of body-odor in women with RPL. To investigate this, we replicated an experiment where we previously observed an impact of body-odors on behavior and psychophysiology (Endevelt-Shapira et al., 2018). We tested 18 RPL women and 18 controls in a single experimental session that included sniffing body-odors, followed by a face recognition task, an empathy task, and an emotional Stroop task, concurrent with Galvanic Skin Response (GSR) recording, once under exposure to Stranger-men body-odor and again under exposure to Blank.

Despite some potentially meaningful group and odorant differences, we failed to observe any significant interaction of group and odorant in this composite set of experiments. In other words, body-odors had the same impact on these measures in both RPL and control women. Although one cannot read too much into such a null effect, it is important to report it when conveying our results, as it confines the extent of difference between these cohorts in their responses to body-odors.

## Discussion

We found that women who experience unexplained RPL have an overwhelming advantage over controls at recognizing their spouse by smell, yet only a slight trend at most towards a general olfactory advantage. Whereas control performance at olfactory spouse identification was on par with two previous reports (Hold and Schleidt, 1977; Mahmut et al., 2019), it was poorer than in a third study, where participants performed on par with the current RPL group (Lundström and Jones-Gotman, 2009). We speculate that these differences across studies reflect methodological differences impacting task difficulty. More specifically, the latter study reapplied body-odor collection-pads for seven straight nights. This likely made for a very concentrated body-odor. In contrast, we reused t-shirts for two nights, making for a less concentrated stimulus, and hence lower overall performance. The question here, however, is not of a comparison across studies, but rather across groups in the same study, where we obviously applied common methods for both groups. In addition to better identification, the body-odor of men strangers smelled differently to women with RPL, and it induced greater hypothalamic activity in RPL versus controls. Finally, all this materialized despite smaller olfactory bulbs and reduced connectivity between the hypothalamus and insula in RPL versus controls. How do these findings combine in relation to our overarching hypothesis of a Bruce-like effect in humans? In the rodent Bruce effect, miscarriage, or pregnancy block, is triggered by the odor of the non-fathering male (Bruce, 1959). For there to be such an olfactory entity as “the odor of the non-fathering male”, the odor of the fathering male must be remembered for comparison. Although there is no reason that this memory be explicit, here we see that women experiencing RPL have a two-fold more accurate explicit memory of their spouse body-odor. This advantage clearly dovetails with the olfactory prerequisites of the *Bruce effect*, and portrays the inability of control women to identify the body-odor of their spouse as a potential adaptation allowing communal living that is prevalent in human societies. Moreover, as noted in the introduction, the Bruce effect entails a brain cascade that culminates in increased release of dopamine from the hypothalamus (Rosser et al., 1989). With this in mind, our finding of increased hypothalamic activation in response to stranger male body-odor in RPL also dovetails with a human Bruce-like effect. This combines with evidence of hypothalamic activation by androgen-like odorants in women but not men (Savic et al., 2001) to imply that body-odors may activate hormonal cascades in humans similar to those at the heart of several social chemosignaling effects characterized in rodents, dovetailing with the growing evidence for social chemosignaling in general human behavior (McClintock et al., 2001; Semin and De Groot, 2013; Luebke and Pause, 2015; de Groot et al., 2017). Moreover, RPL was associated with reduced functional connectivity between the hypothalamus and insula during exposure to stranger men body-odor. In monkeys, connectivity across these regions is related to autonomic regulation (Ongur et al., 1998), and in humans both regions are activated during sexual arousal (Karama et al., 2002). The insula has also been implicated in processing human body-odors related to emotional state (Prehn-Kristensen et al., 2009) and in body-odor-based human kin recognition (Lundstrom et al., 2009). Thus, this pattern further implies altered body-odor related brain processing in RPL within brain substrates relevant to reproduction. Whereas the combination of better spouse identification, increased body-odor-induced hypothalamic activation, and reduced insula-hypothalamus connectivity in RPL can all be woven into a coherent picture consistent with our hypothesis, the finding of reduced olfactory bulb volume in RPL is puzzling. On one side, this result provides yet additional and particularly convincing evidence for an altered olfactory brain profile in RPL. On the other hand, whereas smaller OBs are typically associated with poorer olfaction (Buschhüter et al., 2008; Rombaux et al., 2010), here we observed smaller OBs in the group with better olfactory spouse identification (RPL), and slightly better olfaction overall. A correlation between OB volume and general olfactory performance materialized only within the RPL group, but not in the control group. We note that the relation between OB volume and olfactory performance is not always straight forward (Weiss et al., 2019), and we indeed have no explanation for this relationship in the current data.

We would like to clearly acknowledge some limitations of our study. First, the rodent Bruce effect occurs in pregnancy, yet we cannot test pregnant women. Thus, whether the effects we report increase or diminish during pregnancy remains unknown. Second, and critically, correlation is not causation, and altered behavioral and brain responses to body-odors in RPL, convincing however they may be, do not alone imply that this is causal in the condition. Behavior may differ because of increased motivation in participants with RPL, and brain structure and function may differ because of an independent covarying factor that we failed to identify. Although we could think of several experiments that would investigate causation, these are prevented by ethical considerations. Moreover, any notion of a human Bruce-like effect is further challenged by the underlying neuroanatomy. In rodents, the Bruce effect relies on two neural structures that humans may not possess: The first is the VNO, namely the dedicated sensory epithelia in the rodent nasal passage that is primarily involved in social chemosignaling (Keverne, 1999). Humans may retain a VNO pit on the nasal septum (Trotier et al., 2000), but this structure is considered vestigial in the human nose (Meredith, 2001). Second, the rodent VNO projects to the rodent accessory olfactory bulb (AOB), a structure also reported missing in the human brain (Meisami et al., 1998). It is in the AOB where the odor of the fathering male may be imprinted during mating in rodents (Brennan, 2009). Although humans may lack a VNO and AOB, humans do functionally express vomeronasal receptors in their olfactory epithelium (Rodriguez et al., 2000). Whereas rodent social chemosignaling is primarily attributed to the VNO and AOB, the main epithelium and bulb can also support social chemosignaling (Spehr et al., 2006). Therefore, the missing neural substrates do not altogether preclude a human Bruce-like effect, but do restrict the analogy we make.

With these limitations in mind, we exercised caution in our definitive statements. Our definitive claim, as stated in the title and abstract, is altered perceptual and brain responses to men’s body-odor in RPL. Given this, why do we at all present our results within the context of a proposed Bruce-like effect? First, this hypothesis was genuinely what drove us to investigate RPL in the first place. In other words, altered responses to body-odors in RPL was not a serendipitous finding, but rather a hypothesis-driven finding, and the hypothesis was a Bruce-like effect in humans. Because of the ethical limitations, we opted to start this investigation with a test for circumstantial evidence, and that is what we provided here. This joins with the previously noted evidence ranging from a specific polymorphism in an olfactory receptor gene (OR4C16G>A) that has been associated with RPL (Ryu et al., 2019), to the tradition of protective isolation of women during early pregnancy evident in some tribal societies (Van Gennep, 1960), that together bring us to cautiously air this hypothesis. Such framing provides for a much-needed reorientation of thinking in unexplained RPL. Rather than looking at the uterus and the hormonal environment alone, we looked at the brain, and more specifically at the olfactory system, and identify unique patterns associated with RPL. Indeed, as far as we know, structural and functional patterns of brain organization have not been previously associated with RPL, and this discovery was guided by our overarching hypothesis here. Thus, this framework and set of findings may redirect thinking on this condition, which is currently not well-understood, or managed.

## Materials and Methods

### Participants

Conducting studies in RPL is complicated by the issue of long-term continuous participant availability. This manuscript contains many different experiments that ideally would have been all conducted in the same cohort. This would have allowed optimal relating of one result to another. However, ethical considerations prevent us from conducting any experiments with women who are currently pregnant, or trying to conceive. Because this cohort is often doing exactly that, it is almost impossible to conduct experiments in a given participant over an extended period of time. Instead, participants were available sporadically. To deal with this, we recruited an overall cohort of 40 women with RPL (mean age: 35.06 ± 6.22), and 57 controls (mean age: 34.66 ± 4.24), and then assigned matched pairs based on RPL availability per experiment (Supplementary Table 1). We set the sample sizes based on an initial power analysis based on a previously published study (Lundström and Jones-Gotman, 2009). This implied that at alpha = 0.05 and 80% power we need to test a sample of 29 participants per group. All participants provided written informed consent to procedures approved by the Tel-Hashomer (behavioral studies) or Wolfson hospital (imaging studies) Helsinki Committees. All participants were screened for good general health and no history of psychiatric disease.

### Inclusion/exclusion criteria

Inclusion criteria for RPL was at least two consecutive miscarriages prior to 15 weeks, with an unknown etiology for the pregnancy loss. Inclusion criteria for controls was no history of (known) pregnancy loss or abortions.

### Control matching

For each comparison performed, we matched control participants to RPL participants based on age and number of children. If multiple choices were available, our choice was based on matching the average of the control group as a whole.

## Behavioral studies

### Body-odor collection

Each spouse was provided with a new 100%-cotton white T-shirt for body-odor collection according to standard procedures(Endevelt-Shapira et al., 2018). Briefly, the donors were instructed to wear the shirt for two consecutive nights. The donors were further instructed to avoid eating ingredients that can alter body-odor (fenugreek, asparagus, curry, etc.) prior to body-odor sampling. In addition, during the sampling days, donors were asked not to use soap, shampoo, conditioner or deodorant. Between the two nights, the T-shirts were kept inside a closed glass jar at room temperature, and after the second night, they were further stored at −20°C to prevent bacterial growth.

### Shirt sniffing device (SSD)

On the morning of the experiment shirts were thawed at room temperature inside the jars to avoid humidity condensation. Using sterile scissors, shirts were cut into two longitudinal pieces, such that each half of the shirt contained one axillary area. Each half shirt was then placed inside an SSD – a glass jar covered by a cap with an air filter, inhalation mask and a one-way flap-valve (Supplementary Figure 1). As a blank control, new unworn shirts were also frozen inside a glass jar, and thawed and cut to two halves prior to the experiment.

For each participant, one SSD contained a shirt that originated from her spouse, one SSD contained a shirt that originated from another participant’s spouse (Stranger), and the third contained a Blank. Trials were not time-limited, but on each trial, participants could sniff each jar only once. After sniffing the three SSDs, the participant was asked to identify her spouse from the 3 odors. Participants did not receive feedback as to whether they were right or wrong. Each triangle combination was presented either 4 or 6 times, and percent accuracy was calculated.

### Odorant Identification

We assessed ordinary odorant identification using an established and validated standardized test (University of Pennsylvania Smell Identification Test, UPSIT(Doty et al., 1984)). Although the full UPSIT contains 40 stimuli, the results with a subset of 20 stimuli are highly correlated to the results with 40(Doty et al., 1989). To also verify this in our type of cohort, a subset of 26 RPL participants completed the full 40 stimuli version. We observed a strong fit between the score with 20 and 40 odorants (Spearman Rho = 0.71, p < 0.001). With this in hand, we proceeded to use the 20 stimuli test. Out of 97 participants (40 RPL), 3 were excluded (2 RPL) (>2.7 SD), retaining 94 participants (38 RPL).

### Monomolecules

Each trial contained sequential presentation of three jars (counterbalanced for order), two containing the carrier alone (Propylene glycol) and one containing the carrier with the monomolecule: either androstadienone (AND, androsta-4,16,-dien-3-one, at a final concentration of 0.000025M), androstenone (ANN; 16, (5α)-androsten-3-o, at a final concentration of 0.000025M) or estratetraenol (EST; 1, 3, 5(10), 16-estratetraen-3-ol, at a final concentration of 0.045M). Participants were allowed to take one 2-second long sniff at each odorant presentation, and were then asked to pick out the jar that contained the dissimilar odorant. A subset of participants (same number of RPL and Controls) completed three repetitions per triangle, and the remainder completed 10 repetitions per triangle. The order of presentation of each triplet was randomized, and both the experimenter and participant were blind for condition.

### DMTS threshold test

Olfactory detection thresholds for the alliaceous odor dimethyl trisulfide (DMTS) were determined, using a maximum-likelihood adaptive staircase procedure (MLPEST)(Linschoten et al., 2001), in which a participant was presented with two jars, one blank (containing carrier alone; Propylene Glycol) and one containing a changing concentration of DMTS, and had to choose which contained the odorant (forced choice). Out of 39 participants (19 RPL), 2 were excluded (1 RPL, all > 2.7SD), retaining 37 participants (18 RPL).

### Olfactory Composite Score

The composite score of identification, detection and threshold was calculated using the averaged monomolecule discrimination score, odor identification scores and threshold scores. The two latter scores were normalized to a scale of 0-1 (using the formula (x-min)/(max-min)), and all three scores were then averaged per participant to generate the comprehensive composite score. Out of 97 participants (40 RPL), 1 RPL was excluded (> 2.7 SD) retaining 96 participants (39 RPL). Next, we matched controls to RPL women as described in the “control matching” section, first by matching according to the type of tests these women have a score for, and if multiple options were available, by age and number of children. Three RPL women did not have matched controls by test and age, so we matched controls by the closest type of tests performed and age, and removed the non-matching test from the composite average score of this participant.

### Body-odor ratings

The three body-odors, Spouse, Stranger and Blank, were presented using SSDs to participants in randomized order. Participants were instructed to sniff for 2 seconds and then rate odor on a visual analog scale (VAS) for perceived intensity, pleasantness, sexual attraction and fertility associated with the body-odor. Inter-Stimulus Interval was 30 seconds.

### Behavioral statistics

All data analyses were performed using JASP (JASP Team (2019) Version 0.11.1). Each data-set was first tested for normality of distribution using Shapiro-Wilk test of normality. For normally distributed data, we used the following parametric measures: If only one variable was compared, we used two-tailed t-tests. To test for between-group differences across multiple related variables, we used multivariate analysis of variance (ANOVA) followed by post-hoc two-tailed t-tests. To test for correlations between variables, we used the Pearson correlation coefficient for continuous variables, or Spearman’s Rank Correlation Coefficient if one or both variables were categorical. For abnormally distributed data we used the following nonparametric measures: If only one variable was compared, we used Wilcoxon signed-rank test for one-sample and paired samples, and Mann-Whitney test for independent samples. To test for between-group differences across multiple related variables, we used linear mixed models (using R 3.6.1, packages readxl version 1.3.1, nlme version 3.1-140, car version 3.0-5) followed by the above-mentioned nonparametric tests, and tested for correlations using Spearman’s Rank Correlation Coefficient. Finally, to estimate power, we calculated Cohen’s *d* for parametric measures and rank biserial correlation (RBC) for nonparametric measures. We note that in all cases where we relied on non-parametric tests, we also reported the parametric outcome in parenthesis for comparison. We observe that the non-parametric and parametric approaches yielded nearly identical outcomes in terms of assigning significance to the reported differences.

## Brain imaging

### Participants

A total of 48 participants (24 RPL) were scanned in the fMRI study, and 55 participants (26 RPL) in structural T2-weighted imaging for OB volume and OS depth analysis.

### Data acquisition

MRI Scanning was performed on a 3-Tesla MAGNETOM Trio Siemens scanner. For olfactory bulb (OB) volume and olfactory sulcus (OS) depth, a 32-channel head coil was used. Images were acquired using a T2-weighted turbo spin echo pulse sequence in the coronal plane, covering the anterior and middle segments. Sequence parameters: 35 slices, voxel size: 400 x 400 μm, slice thickness: 1.6 mm, no gap, TE = 85 ms, TR = 7000 ms, flip angle = 120°, 2 averages. Functional data were collected using a 12-channel head matrix coil and T2*-weighted gradient-echo planar imaging sequence, with the following parameters: 450 repetitions, TR = 2000 ms, TE = 25 ms, flip angle = 75°, FOV = 216 × 216 mm^2^, matrix = 72 × 72 mm^2^, voxel size=3×3 mm, slice thickness= 3.7 mm, no gap, 34 transverse slices tilted to the AC-PC plane. Anatomical images for functional overlay were acquired using a 3D T1-weighted magnetization prepared rapid gradient echo (MP-RAGE) sequence at high resolution: 1 x 1 x 1 mm^3^ voxel size, TR = 2300 ms, TE = 2.98 ms, inversion time = 900 ms, and a flip angle = 9°.

#### All imaging data will be made publicly available at https://openneuro.org following acceptance of this manuscript

### Structural brain imaging

Olfactory bulb and olfactory sulcus analysis: We excluded 3 of the 26 RPL participants due to poor resolution (likely motion related) which did not allow a reliable measure of the sulcus or bulb. Next, 23 matching controls were selected based on parity and age of their children, specifically the age of their youngest child, following recent findings regarding the impact of pregnancy on maternal brain anatomy after birth(Kim et al., 2010; Hoekzema et al., 2017; Kim et al., 2018). We used independent sample t-tests to compare OB volume and OS depth between the two groups. OB volume and OS depth were measured according to standard methods(Rombaux et al., 2010). For delineating and measuring the OB we developed software that automatically identifies the region of the OBs, and then allows the user to manually delineate OB boundaries across interpolated images for automated volume estimation. This software was written by co-author LG, has been used before(Weiss et al., 2019), and is available for download at https://gitlab.com/liorg/OlfactoryBulbDelineation. To calculate estimated Total Intracranial Volume (eTIV) we used FreeSurfer version 6.0.0(Buckner et al., 2004), next we normalize the OB by eTIV for 22 participants (MPRAGE for one RPL participants was not obtained, so her matched control was also excluded).

OS depth was measured using ITK-SNAP version 3 (www.itksnap.org). The coronal slice was picked by the Plane of the Posterior Tangent through the eye-balls (PPTE), which in most individuals traverses the anterior-mid segment of the OB. In this slice, a virtual line that was tangent to the inferior border of the orbital and rectus gyri was drawn, and then perpendicular line connecting the above virtual line and the deepest part of the OS was marked. This line represents OS depth(Rombaux et al., 2010). Two independent raters, blind to participant identity and group-allocation, demarcated OB borders and marked OS depth. Next, all the OB volumes and OS depths in which between-rater difference was above 15%, were judged by a third blind review. Subsequent inter-rater correlation in OB volume was r = 0.92, p < 0.001 and in OS depth r = 0.93, < 0.001. Inter-rater agreement was further validated using intraclass correlation coefficient(Tinsley and Weiss, 1975): OB volume: ICC r = 0.96, confidence intervals (CI) 0.925 – 0.981, OS depth: ICC r = 0.967, CI 0.939 – 0.985.

For voxel-based morphometry structural data from 23 RPL and 23 matched-control was processed with FSL 6.0 software. First, the brain extracting tool (BET) was employed to cut the skull from all T1 MRI structural images(Smith, 2002); Next, FSL Automated Segmentation Tool (FAST) was adopted to carry out tissue-type segmentation(Zhang et al., 2001). After careful check for segmentation quality, the segmented GM parietal volume images were aligned to the MNI standard space (MNI152) by applying linear image registration tool (FLIRT) and nonlinear registration (FNIRT) methods. A study-specific template was created by averaging the registered images, to which the native grey matter images were then nonlinearly re-registered. The registered GM parietal volume images were modulated for the contraction/enlargement due to the nonlinear component of the transformation by dividing them by the Jacobian of the warp field. Next, the segmented and modulated images were smoothed with isotropic Gaussian kernels with a standard deviation of sigma = 3 mm. Finally, a voxelwise GLM was applied using permutation-based non-parametric testing, using p < 0.0001, without additional correction for multiple comparisons.

## Functional brain imaging

### Odor stimuli

Twenty-two heterosexual men (mean age 31.1 ± 6.4) donated their body-odor, following the standard procedures described above, with two differences: we used 8 adhesive pads rather than t-shirts, for 4 nights rather than 2. After each night the pads were stored at - 20°C. We used pads rather than t-shirts so as to have smaller and more concentrated odorant sources that would fit into the olfactometer. These pads were cut into small pieces and mixed such that the scanner stimulus contained a mix of 20 men.

### Video clips

An independent group of 12 women rated the emotional arousal, on a scale of 1 to 10, of 30 ~1 min-long scenes. Scenes, all containing human characters, were chosen from 11 commercial films. From these we used the 20 scenes with the highest emotional scores. All scenes were edited into 12-second clips.

### Procedure

To assure no differences related to prior exposure to the film-clips, and to introduce the narrative and emotional context of the stimuli, prior to the scan, participants watched the 20 1-min-long scenes and rated familiarity. Next, a one-hour long fMRI scan was conducted, which included two functional scans, separated by T1 and T2 anatomical scans. Each functional scan started with a 4-sec emotional clip (to be excluded in the analysis) and next 12-sec-long video clips were presented: 20 clips of high emotional content and 20 landscape clips, in alternating order, with an ITI (fixation point) of 8-12 sec. Following each video clip, participants were asked to rate their emotional arousal, on a scale of 1 to 8, with 8 being the highest. Simultaneously with each video clip, body-odor or blank were delivered using a computer-controlled air-dilution olfactometer that embedded the odorant pulse within a constant stream of clean air at 1.5 liter per minute. The two functional scans were identical except for the odor content. The two conditions (blank vs. stranger body-odor) were counterbalanced for order between participants. Participants were aware of the possibility of an odor being delivered, but were not told when. After scanning, participants were asked to report whether there was an odor in each of the sessions, score of “1” = an odor (even if it was noticed only once) and “0” = no odor. First we calculated the differences between body-odor sessions and blank sessions to receive 3 values: 0 (probability: 0.5), −1 (probability: 0.25), 1 (probability: 0.25). Next we used a binomial test to calculate the probability of detection.

Following scanning, participants watched the emotional 12-sec clips again, outside the scanner, and rated them for specific emotions: compassion, happiness, fearfulness, sadness, stress and emotional power, and familiarity prior to the experiment. There were no significant differences between the two groups in any of these ratings (all t(44) < 1.39, all p > 0.17).

### Functional MRI analysis

Preprocessing and functional data analysis were conducted within FSL (FMRIB’s Software Library, www.fmrib.ox.ac.uk/fsl), FEAT (FMRI Expert Analysis Tool) Version 6.00 (Woolrich et al., 2001), and MATLAB R2018a (MathWorks, Inc.).

### Preprocessing pipeline

Two participants were excluded from the analysis: one RPL woman due misunderstanding of the task (she did not rate the emotional arousal of the landscape clips), and one control participant due to miscarriage that was reported only after the scanning. Thus, a total of 46 participants, 23 in each group were submitted to further analysis. The first 8 volumes (4-s clip) in each run were discarded to allow the MR signal to reach steady-state equilibrium. The structural brain images were skull-stripped using the FSL brain extraction tool(Smith, 2002). Functional images were corrected for slice-timing and head motion using 6 rigid-body transformations with FSL MCFLIRT(Jenkinson et al., 2002). Within each run, functional images were spatially normalized to the individual’s anatomy and co-registered to the MNI 152 T1 template, using a combination of affine and non-linear registrations. Images were spatially smoothed with an 8 mm Gaussian kernel, and a high-pass filter (cut off = 128 sec) was incorporated into the GLM to correct for scanner drift.

### Voxel-wise analyses

On the first level, all 3 EV regressors were modeled by a stick function convolved with a double gamma function. The regressors included: All video-clips with no weightings (and their temporal derivative), video-clips weighted with emotional arousal ratings (demean), reaction time (orthogonal to the other 2 regressors). This model also included a regressor of no interest for each volume, with > 0.9 mm framewise displacement (using FSL motion outliers). At the second level, the parametric estimation of the weighted-clips was a contrast of stranger body-odor>blank using fixed-effects. Finally, for between group analysis we used small-volume correction of the hypothalamus (defined by Neurosynth), and FLAME1 (FMRIB’s Local Analysis of Mixed Effects), with cluster-size correction z>2.3 (p < 0.01).

### Functional connectivity

We used psychophysiological interaction (PPI) analysis(Friston et al., 1997) to measure changes in functional connectivity modulated by emotional arousal. We conducted whole-brain PPI tests, reflecting greater correlation with hypothalamic (seed ROI) time series (physiological regressor) for all emotionally-weighted-movie-clips > fixation (psychological regressor). At the second level, the parametric estimation of the weighted-clips was a contrast of body-odor>blank using fixed-effects. Finally, for between group analysis we used FLAME1, with cluster-size correction z>3.1 (p<0.001).

## Acknowledgments

This study was funded by a European Research Council AdG. grant #670798 (SocioSmell) awarded to N.S. General work in the Sobel lab is funded by the Rob and Cheryl McEwen Fund for Brain Research. Dr. E. Furman-Haran holds the Calin and Elaine Rovinescu Research Fellow Chair for Brain Research

## Author contributions

Designed experiments: LR; RW; TW; HC; NS. Ran behavioral studies: LR; RW; IR; IF; SS; NR; YH; LP; YES; EM. Analyzed behavioral data: LR; RW; TW; IR; LG; YES; EM; NS. Ran imaging studies: LR; RW; TW; SS; LG; EFH. Analyzed imaging data: LR; RW; TW; SS; LG. Wrote the first draft: LR; RW; TW; NS. Commented/edited manuscript: All authors.

